# Co-translational targeting of transcripts to endosomes

**DOI:** 10.1101/2020.07.17.208652

**Authors:** Doris Popovic, Wilco Nijenhuis, Lukas C. Kapitein, Lucas Pelkmans

## Abstract

Asymmetric localization and translation of mRNAs is used by single cells to sense their environment and integrate extrinsic cues with the appropriate cellular response. Here we investigate the extent to which endosomes impact subcellular patterning of transcripts and provide a platform for localized translation. Using image-based transcriptomics, indirect immunofluorescence, and RNAseq of isolated organelles, we discover mRNAs that associate with early endosomes in a translation-dependent and -independent manner. We explore this in more detail for the mRNA of a major endosomal tethering factor and fusogen, Early Endosomal Antigen 1, *EEA1*, which localizes to early endosomes in a puromycin-sensitive manner. By reconstituting EEA1 knock-out cells with either the coding sequence or 3’UTR of EEA1, we show that the coding region is sufficient for endosomal localization of mRNA. Finally, we use quantitative proteomics to discover proteins associated with *EEA1* mRNA and identify CSRP1 as a factor that controls EEA1 translational efficiency. Our findings reveal that multiple transcripts associate with early endosomes in a translation-dependent manner and identify mRNA-binding proteins that may participate in controlling endosome-localized translation.

## Introduction

Compartmentalized localization of mRNAs has been recognized as an important regulator of fundamental processes that govern cell polarity, cell fate, cell movement, and differentiation, including early embryonic patterning and asymmetric cell division^1^. There are several known mechanisms that enable non-random distribution of transcripts, such as localized protection of mRNA from degradation, anchoring and directed movement of mRNA along the microtubules, and more recently discovered – trafficking of mRNA and ribonucleoparticles on late endosomes^2–4^ and mitochondria^5–7^. These mechanisms are mediated by a number of RNA-binding proteins that recognize secondary structures within 3’UTR regions of mRNAs and influence nuclear export, cytoplasmic movement, and accessibility of mRNA to both ribosomes and mRNA degradation machinery. For some transcripts, these mechanisms have been explored in detail, such as for the mRNA that encodes beta actin (ACTB) in various mammalian cells^8^, and for the mRNA that encodes the transcriptional regulatory protein Ash1 in yeast^9,10^. Furthermore, in *D. melanogaster*, numerous mRNAs have been shown to localize at specific sites during embryogenesis, and such patterned localization is essential for proper development and formation of the anterior-posterior axis in early *D. melanogaster* embryos^11–13^. In more specialized cells, such as neurons, anchoring of mRNA on late endosomes enables delivery of mRNA along the axon, thereby controlling appropriate context-dependent spatial organisation of the proteome and metabolites^2^. Additionally, localized translation of functionally related mRNAs can drive efficient assembly or larger protein complexes that are immediately engaged in essential cellular processes, such as nuclear pore proteins during nuclear pore assembly upon cell division^14^.

We previously developed an approach that we termed image-based transcriptomics, which utilizes a sensitive branched DNA single-molecule fluorescence *in situ* hybridization (bDNA sm-FISH) technique to visualize and detect single transcripts in thousands of single cells and extract numerous features reflecting their subcellular spatial patterning in high-throughput^15,16^. This enables unbiased systems-level analysis of the types of intracellular transcript patterning and their correlation with other cellular properties (such as cell size, position, microenvironment, and neighbour activity) at a large scale^17^. This has shown that cytoplasmic transcript abundance can for most genes be largely explained by the phenotypic and microenvironmental state of single cells^18^, and that such features, in combination with quantifying mRNA subcellular patterning, can explain mRNA-to-protein ratios across populations of single human cells for different types of gene expression profiles^19^. This revealed that the cell-to-cell variability in transcript abundance and mRNA translation efficiency emerges as a consequence of the self-organisation of adherent mammalian cells into populations, whereby each individual cell adapts its phenotype to the locally created multicellular microenvironment.

Here, we combined image-based transcriptomics with indirect immunofluorescence imaging to quantify the patterning of transcripts of 328 different genes encoding for endocytic and endomembrane machinery, using our previously published library of smFISH probes^15^. This allowed us to directly correlate spatial patterns of transcripts with the position of early and late endosomes for both translationally active and ribosome-dislocated transcripts. Together with RNAseq of an affinity-isolated early endosomal cell lysate fraction, this revealed that transcripts of a subset of genes encoding for endocytic machinery, signal transduction factors, and cytoskeleton regulators associate with early endosomes in a translation-dependent manner. For one of such transcripts, the mRNA of Early Endosomal Antigen 1 (*EEA1*), a well-characterized molecular tether that increases homotypic fusion of early endosomes ^20,21^, we show that its protein-coding sequence is sufficient for endosomal targeting, and does not depend on the protein nascent chain binding to endosomal membranes. Using quantitative mass spectrometry, we identified *EEA1* transcript-associated proteins and show that one of these proteins, CSRP1, acts as a negative regulator of *EEA1* mRNA translation. Our findings establish endosomes as mRNA localisation sites for regulated translation by additional RNA-binding factors. Given extensive contacts between endosomes and the endoplasmic reticulum^22,23^, these findings may reflect mechanisms by which the translation of transmembrane or secreted proteins and cytoplasmic proteins with functional roles in the endo-lysosomal pathway is coordinated.

## Results

### Image-based transcriptomics reveals endosome-localized transcripts

To study whether transcripts display endosomal patterning, we combined bDNA smFISH with indirect immunofluorescence (Fig 1A). smFISH was performed against 328 genes, encoding for various endosomal and endomembrane regulators, including those that are translated on the ER, such as transmembrane growth factor receptor proteins, as well as cytoplasmic proteins that associate with endosomes and are regulators of signaling and endocytosis (Fig 1B). We imaged the localization of transcripts in cells undergoing active translation and in cells in which translation was inhibited by the addition of puromycin, which dislocates mRNAs from ribosomes. The obtained transcript counts were highly reproducible and did not alter during puromycin treatment (Fig S1A). Using indirect immunofluorescence, we stained both early (EEA1-positive) and late (LAMP1-positive) endosomes. Next, we measured single pixel-correlations between smFISH signals and the two endosomal stains in both conditions and quantified the changes in spatial features of mRNA molecules in single cells, across thousands of cells (Fig 1A). From these features, we determined 5 types of spatial patterns that transcripts adopt in individual cells, namely perinuclear non-clustered (class 1), peripheral (class 2), peripherally clustered (class 3), peripherally non-clustered (class 4) and spread-out (class 5), as described previously^15^. This revealed that transcripts of functionally related genes tend to display similar spatial patterns (Fig 1C). For instance, we observed that genes that encode for subunits of a common protein complex or regulatory signaling pathway frequently occupy the same cluster, such as Mitogen-activated Protein Kinases (MAPK) and its regulators (cluster 3), autophagy, peroxisomal and Wnt pathway genes (cluster 5), late endosomal genes (cluster 6), early and recycling endosome-associated RAB GTPases, regulators of clathrin-mediated endocytosis and subsets of components that belong to endosomal sorting complexes required for transport (ESCRT) (cluster 7), SMAD genes and recycling RAB11 dependent machinery (cluster 8) (Fig S1D)^15^. This underscores the notion that the subcellular localization of many transcripts is non-random, as reported previously by us and others ^15,24^.

**Figure 1.**
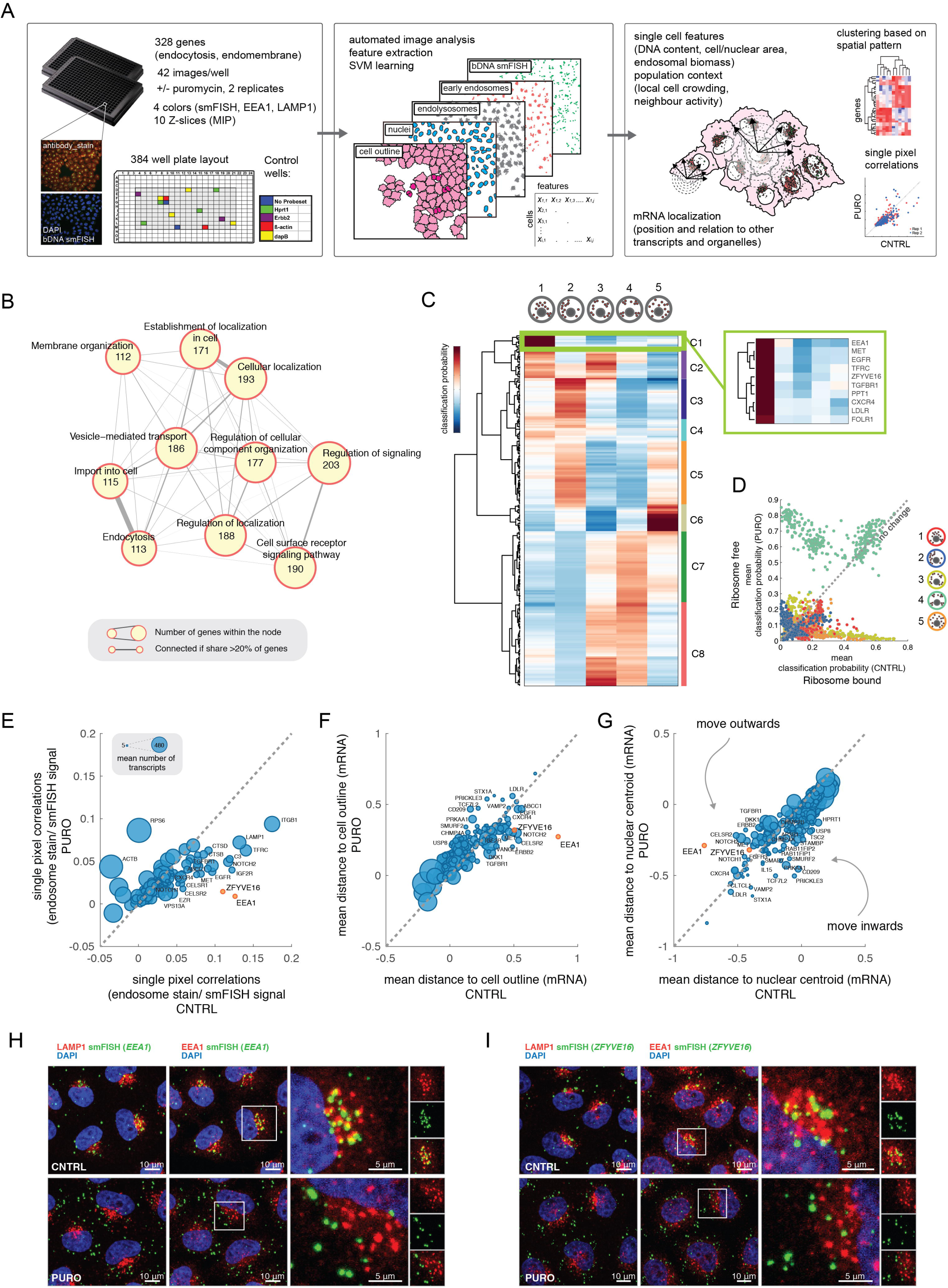
Image-based Screen for mRNAs that Localize on Endosomes. (A) Scheme of the workflow, left to right: number of inspected genes in two different conditions, detection and quantification of mRNA abundance and localization, extraction of cellular features, and data clean-up (badly segmented cells and border cells cleaned by SVM (Supervised machine learning). Puromycin treatment was done for 1hr (1 µg/ml). MIP (maximum intensity projection). (B) Gene Ontology enrichment network of 328 genes used for the image-based screen, genes were grouped based on Biological Process. Width of the interaction represents number of shared genes between nodes. Size of the node is based on the number of genes that belong to the node. Nodes are connected only if they share more than 20% of genes. GO annotation was performed using gProfiler algorithm. (C) Hierarchical clustering of genes based on their spatial class probabilities to belong to one of the 5 spatial patterns. Cluster 1 is highlighted as containing candidates most prominently localized on endosomes. (D) Correlation plot of classification probabilities in control (ribosome bound) against puromycin treated conditions (ribosome free) for all tested genes. (E) Correlation of single pixel correlations with the endosomal stains in control condition against puromycin treatment condition. EEA1 and ZFYVE16 are highlighted as strongest candidates. Genes with less than 5 spots per cell were excluded from analysis. (F) Correlation of mean mRNA distance from the cellular outline in control condition against puromycin treatment condition. (G) Correlation of mean mRNA distance from the nuclear centroid in control condition against puromycin treatment condition. (H) Example images of *EEA1* mRNA, EEA1 protein and LAMP1 protein localization in control and puromycin condition. (I) Example images of *ZFYVE16* mRNA, EEA1 protein and LAMP1 protein localization in control and puromycin condition.

We next asked which mRNAs co-localized with endosomal markers, which predominantly localize in the perinuclear region of the cell. This is similar to the type 1 spatial patterning of transcripts (Fig 1C), which contain genes that encode for a specific subset of ER-translated growth factor receptor proteins (eg. *EGFR, TGFR, TFRC, MET, CXCR4, LDLR, FOLR1*), but also genes that encode for cytosolic proteins without signal sequence, such as *EEA1* and Zinc Finger Containing protein *ZFYVE16*. Interestingly, transcripts of genes encoding for other transmembrane receptors, such as *NOTCH* or the Cadherin family of receptors displayed a different more peripheral spatial patterning (class 5) (Fig S1D). This indicates that ER-localized translation of transcripts may be spatially organized depending on the type of transcript being translated. When we compared the spatial patterning of transcripts between control and puromycin-treated cells, we observed many pattern changes (Fig 1D). This was also evident in the single-pixel correlations between smFISH and endosomal protein stain signal (Fig 1E) as well as in the mean distances of single mRNAs from the cellular outline (Fig 1F) or the nuclear centroid (Fig 1G). In particular the transcripts of *EEA1* and *ZFYVE16*, that belonged to cluster 1, showed a clear decrease in correlation (Fig 1H). This effect was to some extent also observed for specific transcripts encoding transmembrane proteins that have a function in the endosomal pathway (*LAMP1,TFRC*). In addition, we observed for transcripts of *EEA1* also a clear outwards translocation upon addition of puromycin (Fig 1I). Taken together, this indicates that for many transcripts, their subcellular patterning is different depending on their ribosome-associated translational state. These specific patterns may reflect localization to endosomes or other organelles depending on the cellular function of the gene, and may even apply for genes that are known to be translated by ER-bound ribosomes, suggesting that the heterogeneity in patterning of ER-associated transcripts^25,26,15^, has a biological function that may emerge from physical contacts between the ER and other organelles^23,27^. Understanding the underlying heterogeneity of ER-associated transcript patterning will require further characterization, and we here focus on studying transcripts of cytosolic proteins associating with endosomes.

### RNA sequencing of isolated endosomes reveals endosome-associated transcripts

To confirm our findings obtained with image-based transcriptomics, and to identify additional transcripts that associate with the endosomal compartment in a translation-dependent manner, we used RNA sequencing as an orthogonal approach. Using magnetic beads coupled to either an antibody against EEA1 protein as an early endosomal marker, or an antibody against CKAP4, which is an ER-sheet resident protein, we immunoprecipitated respectively fractions of endosomes or the ER from lysates obtained from both untreated and puromycin-treated cells (Fig 2A, Fig S2A, Fig S2B). Subjecting these fractions to Next Generation Sequencing identified numerous specific transcripts in the different fractions with high reproducibility among the biological replicates (Fig S2C, S2D). Importantly, this identified transcripts of both *EEA1* and *ZFYVE16* in the endosomal fractions of untreated cells in all 4 replicates, but not in endosomal fractions of puromycin-treated cells or in ER fractions (Fig 2B and 2C), confirming our image-based transcriptomics results. Subsequent Gene Ontology (GO) enrichment analysis revealed that the endosomal fraction of untreated cells, as well as the endosomal fraction of puromycin-treated cells compared to their respective ER fractions (Fig 2B) comprised primarily of transcripts encoding for cytosolic proteins involved in endocytic trafficking and signaling, as well as in the organization of the actin cytoskeleton, but no transmembrane receptors. When we inspected the top-100 transcripts significantly enriched in the endosomal fraction compared to the ER fraction in a puromycin sensitive-manner, we identified, besides transcripts of *EEA1* and *ZFYVE16*, several transcripts encoding components of the WASH complex (*FAM21C, FAM21A, FKBP15, KIAA1033*), known to mediate actin cytoskeleton dynamics. Furthermore, we identified transcripts encoding *SYNE2*, a component of the LINC (Linker of Nucleoskeleton and Cytoskeleton) complex that bridges organelles and the actin cytoskeleton, transcripts encoding regulators of clathrin-mediated endocytosis (*ITSN2, SYNRG*), an ESCRT complex phosphatase (*PTPN23*), a regulator of EGFR endocytosis and MAPK signaling (*NISCH*), Ribosomal Protein S6 Kinase C1 (*RPS6KC1*), as well as a kinase involved in sphingosine-1-phosphate signaling (Fig 2C-D).

**Figure 2.**
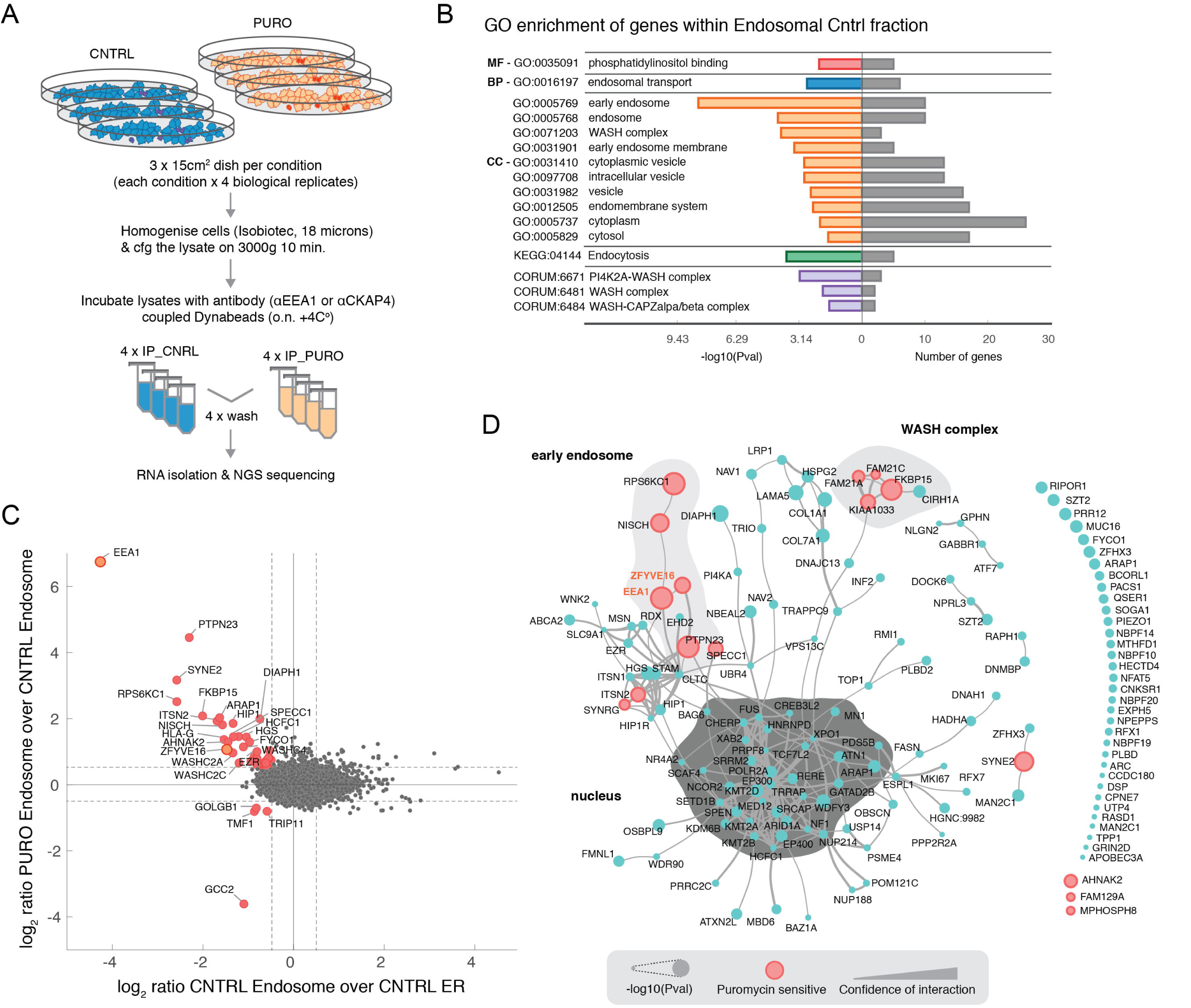
RNA sequencing of Purified Endosomes Reveals Associated Transcripts. (A) Scheme of the workflow for the sample preparation for the RNAseq of purified intracellular organelle fractions. Puromycin treatment was done for 1 hr (1 µg/ml). (B) Gene Ontology enrichment of genes detected to be significantly present in endosomal fraction (all 4 replicates). MF – Molecular Function; BP – Biological Process; CC – Cellular Component. (C) Correlation plot of log_2_ transformed ratios of control endosomal fraction over control ER fraction against log_2_ transformed ratios of puromycin treated endosomal fraction over control endosomal fraction. Red highlighted genes in the upper left corner represent those significantly enriched within the endosomal fraction, that are at the same time sensitive to puromycin. (D) Top 100 genes significantly enriched within the endosomal fraction (as compared to the ER fraction, p value cutoff 0.05). Confidence of interaction was obtained from STRING. Genes with no interaction reported are listed on the right side of the network. Genes sensitive to the Puromycin treatment are highlighted in red.

As transcripts of *EEA1* were highest enriched in the endosomal fraction and displayed clear colocalization with endosomes in a puromycin-dependent manner, we decided to focus on this gene to further investigate the mechanism and function of endosomal localization of transcripts. The EEA1 protein forms an extended coil-coiled homodimer with an N-terminal zinc finger domain that binds to the early endosomal GTPase RAB5 and a C-terminal FYVE domain that binds to PI3P on early endosomes, thereby acting as a tethering molecule that increases the efficiency of homotypic early endosome fusion^17,18,24^. As such, EEA1 plays an important role in controlling the rate and degradation dynamics of internalized EGFR receptor and the cellular capacity to process and decode information about growth factor concentration in the extracellular environment^29^. We next asked whether endosomal localization of *EEA1* mRNA is a conserved phenomenon across 7 different human cancer-derived cell lines (HeLa, CaCo2, HepG2, HCT, U2OS, A549, A431), two non-transformed human cell lines with a stable genome (RPE1 and MRC5), and a Green African monkey cancer cell line (Cos7). Across all cell lines inspected, *EEA1* mRNA colocalized with the EEA1 protein on endosomes in control conditions, and was displaced from endosomes upon puromycin treatment, as evident by a significant decrease in measured single pixel correlations, and changes in distances of single mRNA molecules from the center of the nucleus and cellular outline (Fig S3A). When we calculated the spatial class probabilities across all the single cells in two different conditions, puromycin treatment resulted in a clear change in mRNA localization from perinuclear to spread out, in virtually all inspected cell lines (Fig S3B, S3C). This conservation of co-translational *EEA1* mRNA targeting to endosomes across various mammalian cell types suggests that it serves an important regulatory role. Localized translation of *EEA1* mRNA on the surface of endosomes may for instance control the number of early endosomes, and consequently influence cargo degradation and EGFR signaling, on time scales that can be shorter than those involving transcriptional control.

### *EEA1* transcripts are anchored to early RAB5-positive endosomes

We next asked whether colocalization of *EEA1* mRNA to early endosomes involved anchoring to the endosomal membrane. To test that, we depolymerized microtubules by treating cells with a high concentration of nocodazole, which results in scattering of early endosomes throughout the cytoplasm. This also resulted in the concomitant scattering of *EEA1* transcripts, but they remained co-localized to endosomes, in contrast to the effect of puromycin (Fig 3A). While both treatments changed the mRNA patterning from a more perinuclear to a more spread-out localization (Fig 3B), their effects were different. Upon nocodazole treatment, transcripts remained clustered in their more peripheral locations (class 4), while they were less clustered (class 3) and more randomly distributed (class 5) upon puromycin treatment. Importantly, while both treatments did not or only minimally affect *EEA1* transcript counts (Fig. 3C) and EEA1 protein content (Fig. 3D), the transcript distance to the nuclear centroid increased more in the presence of puromycin compared to nocodazole (Fig. 3E), while the single-pixel correlations between *EEA1* mRNA and EEA1 protein stains decreased more in the presence of puromycin compared to nocodazole (Fig. 3F).

**Figure 3.**
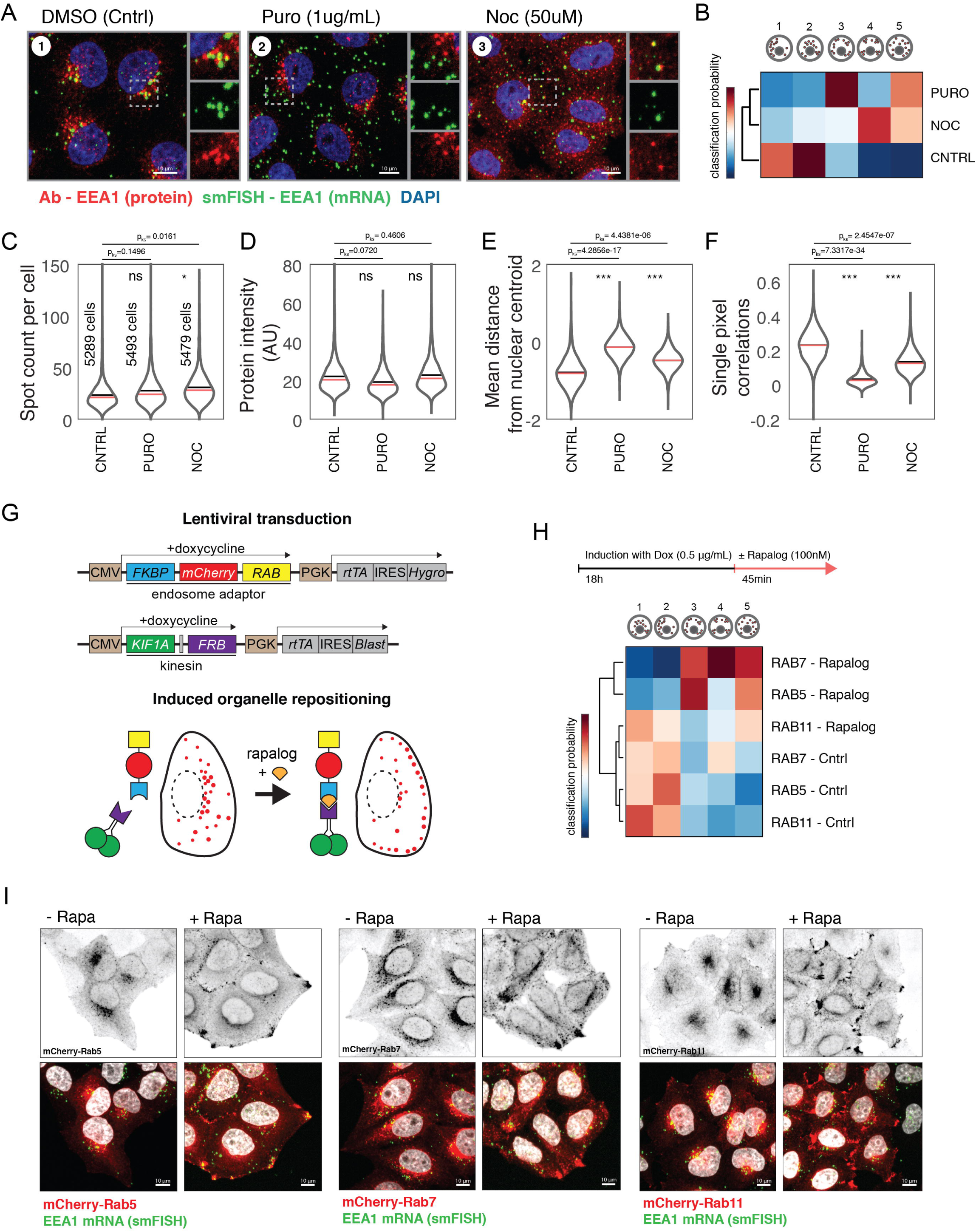
*EEA1* mRNA Associates with the Early RAB5 endosomes. (A) Representative images of EEA1 mRNA colocalization with the early endosomes in Control, Puromycin and Nocodazole treated conditions. Puromycin and Nocodazole treatment were performed for 1hr. (B) Hierarchical clustering of conditions based on the spatial class probabilities for *EEA1* mRNA to belong to one of the 5 spatial pattern classes. (C) mRNA count of EEA1 mRNA across three conditions as described in (A). p values for KS statistics were obtained by bootstrapping 100 times single cells from the total pool and comparing the distributions of Control against treated condition. (D) Protein levels of EEA1, across three conditions as described in (A). p values for KS statistics obtained same as in (B). (E) Single pixel correlations of sm-FISH signal and EEA1 protein stain, across three conditions as described in (A). p values for KS statistics obtained same as in (B). (F) Mean mRNA distance from the nuclear centroid, across three conditions as described in (A). p values for KS statistics obtained same as in (B). (G) Scheme of organelle-repositioning system: a panel of cell lines was generated, stably expressing KIF1A-FRB and FKBP-mCherry-RAB5, 7 or 11 from a doxycycline-inducible promoter. Heterodimerization of FKBP-FRB upon rapalog addition induces peripheral enrichment of specific subsets of endosomes (red). (H) Upper arrow represents timeline of induction of expression (Dox) and dimerization (Rapalog) for each of the inspected cell line, expressing different RAB GTPase. Lower clustergram represents hierarchical clustering of the conditions based on the spatial class probabilities of EEA1 mRNA to belong to a particular spatial class pattern. (I) Representative images of HeLa cells expressing RAB5, RAB7 or RAB11, respectively, in non-translocated condition (-Rapa), and translocated condition (+Rapa).

To identify the type of endosomes that carry EEA1 mRNA, we sought to displace specific classes of endosomes from the crowded perinuclear region to the cellular periphery. For this, we made use of a chemical heterodimerization system to induce coupling of kinesin-3-FRB to FKBP-mCherry-tagged endosomal adaptor proteins upon the addition of the small molecule rapalog, thereby stimulating acute and precise translocation of specific endosomes towards the cell periphery^30^. For this purpose, we generated a panel of cell lines stably expressing KIF1A(1-365)-FRB and either RAB5 (early endosomes), RAB7 (late endosomes), or RAB11 (recycling endosomes) tagged with FKBP-mCherry (Fig. 3G), all expressed from a doxycycline-sensitive promoter. We found that endosomes became peripherally enriched in all mCherry-positive cells after 45 min rapalog treatment, at which point we performed smFISH and quantified the mRNA spatial features across thousands of single cells. This revealed that translocation of early RAB5-positive endosomes resulted in the most severe re-localization of *EEA1* mRNA to the cellular periphery, where it colocalized with RAB5. Translocation of RAB7-positive endosomes induced such a change to a lesser extent, while RAB11 translocation did not affect EEA1 mRNA localization (Fig 3H, I). This shows that *EEA1* transcripts primarily associate with early endosomes and suggests that they can be transported to specific destinations in the cell through the directional movement of endosomes. Interestingly, mRNA transport via endosomes has previously been described in the context of late endosomes, where ribonucleoparticles are tethered to the endosomal membrane via Annexin A11^4^. In *Ustilago maydis*, trafficking of mRNA on early endosomes was reported to be depended on RNA binding protein Rrm4^3^.

### cDNA but not 3’UTR is sufficient for endosomal recruitment of EEA1 mRNA

To explore the mechanism of EEA1 mRNA tethering to endosomes, we next developed assays for reconstitution in *EEA1* knock-out cells. One possibility is that the puromycin-sensitive *EEA1* mRNA localization to RAB5-positive endosomes is mediated by the nascent polypeptide of EEA1 emerging from the ribosome as the transcript is translated, since its ability to bind to RAB5 is mediated by a Zinc finger domain at its N-terminus. In this scenario, the N-terminal domain would immediately bind to RAB5 on endosomal membranes upon exit from the ribosome. To address this, we generated an *EEA1* knock-out (KO) cell line using CRISPR/Cas9, and reintroduced different variants of *EEA1*-encoding plasmids by stable integration within unique LoxP sites. Specifically, we rescued cells with either the full cDNA sequence of *EEA1* tagged with EGFP, or one of three different mutant forms tagged with EGFP deficient in i) binding to RAB5 by having two point mutations in the N-terminal zinc finger (E39A/F41A), ii) deficient in binding to PI3P by introducing a stop codon before (at position 1349), or iii) having a point mutation (H1373A) in the C-terminal FYVE domain, which abolishes PI3P interaction. Additionally, we added the 3’UTR of *EEA1* to the coding sequence of EGFP and reintroduced it into the KO HeLa cell line, as 3’UTR regions have well-known roles in in mRNA localization in other genes (Fig S3D). We then induced the expression of these variants for 8 hours, and quantified abundance and localization of the transcripts using smFISH against the EGFP sequence. This revealed that transcripts with the cDNA sequence of EEA1, including those with mutations, displayed spatial patterns typical of endosomal association (closer to the nuclear periphery, further away from cell outline), even though the mutated forms of the EEA1 protein did not localize to endosomes (Fig. S3D, E). In contrast, transcripts containing only the 3’UTR sequence of *EEA1* did not display a spatial pattern typical of endosomal association but were distributed throughout the cell. In addition, their abundance was 30-fold higher (Fig. S3D, E). This indicates that although localization of *EEA1* transcripts is puromycin-sensitive, and thus likely involves association to ribosomes, it does not depend on the ability of the nascent or full-length protein to be able to bind to RAB5 or PI3P. This ability appears to be conferred by the coding sequence of *EEA1* mRNA, which may be coupled to an increased turnover of the transcripts.

### SILAC-based proteomics of EEA1 mRNA reveals RNA-binding proteins

Co-translational targeting of transcripts to the endoplasmic reticulum membrane (ER) is well characterized for transcripts that encode for transmembrane or secreted proteins. This involves classical signal peptide-dependent targeting, which relies on the signal recognition particle (SRP) to bind to the signal peptide in the nascent protein, triggering a conformational change that enables binding to the SRP receptor on the ER membrane, which is associated with the translocon^31–33^. Recently, additional mechanisms of targeting transcripts to the ER that depend on other ribosome- and mRNA-associated factors have also been discovered^34–36^. While still relying on ribosome association, these mechanisms do not depend on some property in the nascent protein. Such ribosome association-dependent localization of transcripts has also been described for organelles other than the ER, such as mitochondria^5,6,7^.

To identify mRNA-associated factors that might be involved in the case of *EEA1* transcript targeting to endosomes, we developed an assay for the specific isolation of endogenous *EEA1* transcripts together with its associated proteome. To this end, we used magnetic beads coupled to oligonucleotides complementary to the sequence of the *EEA1* transcript, commonly used for single-cell RNA sequencing purposes. We then applied quantitative proteomics using Stable Isotope Labeling by Amino Acids in Cell Culture (SILAC) to identify proteins associated with isolated *EEA1* mRNA from HeLa cell lysates. Magnetic beads coupled to oligonucleotides against *EGFP* mRNA (which was not expressed in these cells) were used as background control (Fig 4A., Fig S4A). In a parallel experiment, we applied to same approach to identify proteins associated with *ACTB* mRNA (Fig S4B). Subsequent analysis of proteins associated with both *EEA1* or *ACTB* transcripts showed, as expected, an enrichment for RNA-binding proteins and common factors involved in mRNA translation (Fig 4B and Fig S4B), raising confidence in the approach. Analyzing the proteins specifically bound to *EEA1* mRNA (Fig. 4B) revealed several that have previously been functionally implicated in modulating signaling, endocytosis, and the actin cytoskeleton, including Ankyrin Repeat Domain 13A (ANKRD13A), which binds to ligand-activated, ubiquitinated EGFR at the cell membrane to mediate its endocytosis and downregulation; Protein Kinase N2 (PKN2), a Rho/Rac effector protein; Neuronal Tyrosine-Phosphorylated Phosphoinositide-3-Kinase Adaptor 2 (NYAP2), which activates PI-3 kinase and recruits the WAVE1 complex to trigger actin nucleation; and Cysteine And Glycine Rich Protein 1 (CSRP1), a LIM domain containing protein^37^ that interacts with actin and the actin bundling protein α-actinin, thereby facilitating actin bundling^38^, and may control integrin-dependent cell migration^39^. The functional relationship between these cellular activities and the well-established role of EEA1 in endocytosis prompted us to test their importance in *EEA1* mRNA localization and translation.

**Figure 4.**
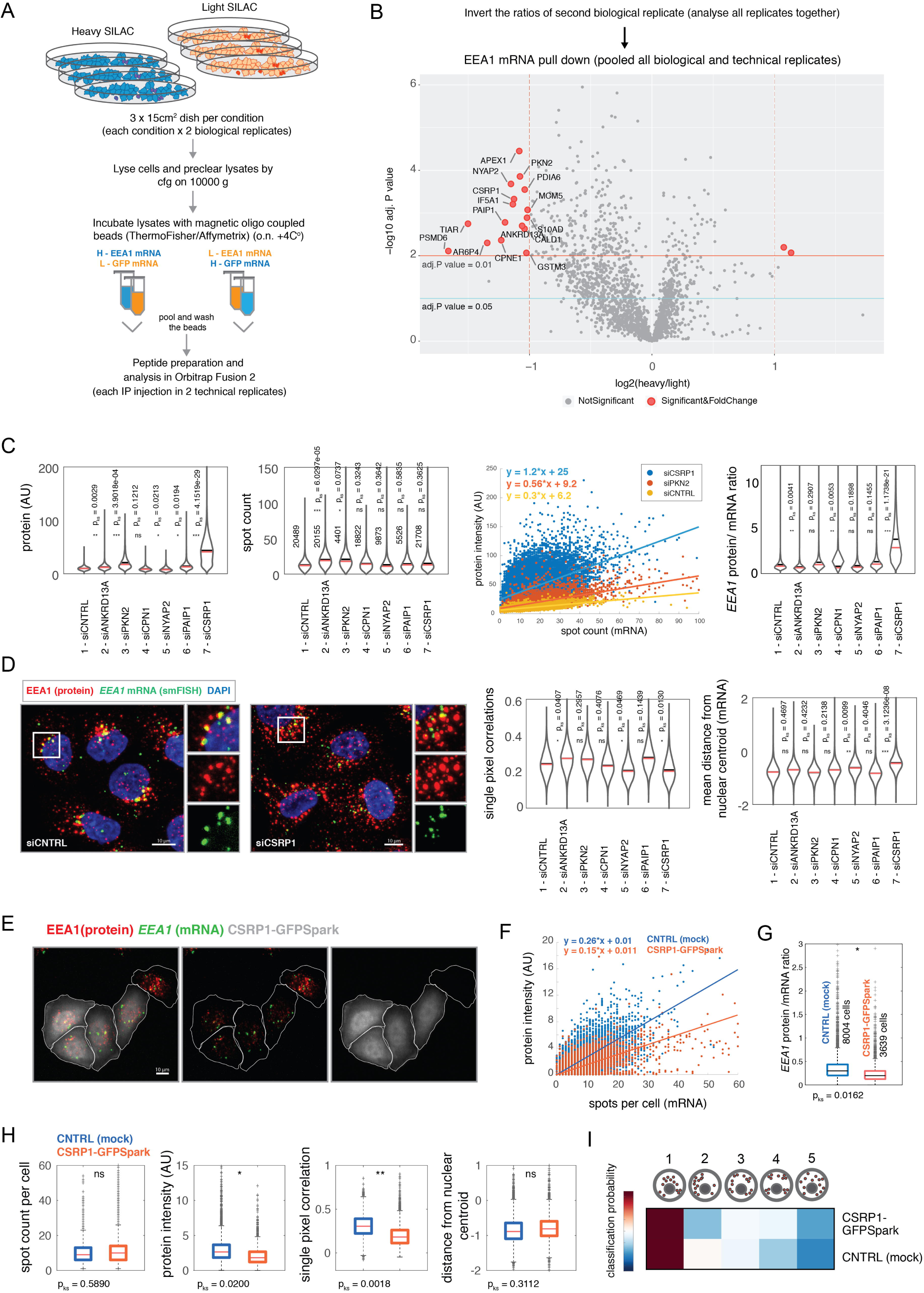
SILAC Proteomics Reveals Interactors of *EEA1* mRNA. (A) Scheme of the workflow for the specific isolation of the *EEA1* mRNA, and SILAC labeling of the HeLa cells in culture. Two biological replicates were samples with reverse labeling. (B) Analysis of two biological and two technical replicates. Proteins significantly enriched bound to the EEA1 mRNA are highlighted in red (p<0.01, log_2_<-1), left side of the volcano plot. (C) Left: mRNA count and EEA1 protein levels in control siRNA and siRNA conditions of selected mRNA binding proteins. Right: Linear regression plots using mRNA count as a predictor for the protein quantity (EEA1) and protein to mRNA ratios in siCNTR or siRNA against RNA binding protein treated cells. P values for KS statistics were calculated as in (C). (D) Left: Representative images of siCNTRL cells and siCSRP1 cells. Right: Single pixel correlations of sm-FISH signal and EEA1 protein stain in control siRNA and siRNA conditions of selected mRNA binding proteins; All p values for the KS statistics were calculated by bootstrapping 100 times population of single cells and comparing the distributions of the siCNTRL treated cells against the gene targeted siRNA treated cells; Mean mRNA distance from the nuclear centroid in control siRNA and siRNA conditions of selected mRNA binding proteins. (E) Representative images of HeLa cells transfected with the CSRP1-Spark-GFP plasmid. (F) Linear regression using mRNA count as a predictor for the protein quantity (EEA1) in transfected cells (red) and non-transfected cells (blue). (G) Protein to mRNA ratios in transfected (red) and non-transfected (blue) cells. (H) Left to right: *EEA1* mRNA count, protein quantity, single pixel correlations between sm-FISH and protein stain signal and mean mRNA distance from the nuclear centroid in transfected (red) and non-transfected (blue) cells. P values for KS statistics were calculated by bootstrapping 100 times cells from both populations and comparing the distributions of measured feature. (I) Heatmap of mean spatial class probabilities based on *EEA1* mRNA spatial localization in CSRP1-GFPSpark transfected and non-transfected cells.

### CSRP1 levels negatively correlate with the EEA1 protein levels

To test a role of the identified *EEA1* mRNA-binding proteins in the localization and translation of *EEA1* transcripts, we performed siRNA-mediated gene silencing. 72 hours after siRNA transfection, we fixed cells and inspected the localization and abundance of both *EEA1* mRNA and protein. Among the genes tested, silencing of particularly *CSRP1* led to a strong, 5-fold increase in EEA1 protein abundance (Fig. 4C), while mRNA abundance was unchanged (Fig. 4C). Plotting the ratio between *EEA1* protein and mRNA across a large number of single cells revealed that while scaling between these two abundances was largely linear in both control cells and CSRP1-silenced cells, the slope was ∼4-fold steeper in the latter cells (Fig. 4C). Consistently, when we calculated the protein-to-mRNA ratios across all single cells, we observed that in CSRP1-silenced cells ratios significantly increased as compared to the siRNA control cells (Fig. 4C). We also observed that *EEA1* mRNA was more peripherally located and single-pixel correlations between *EEA1* mRNA and EEA1 protein stains were somewhat lower in CSRP1-silenced cells compared to siRNA control cells (Fig. 4D, Fig. S4E).

We then asked whether an increase in CSRP1 levels would induce the opposite effect. Overexpressing GFP-tagged CSRP1 resulted in a decreased EEA1 protein abundance, without affecting *EEA1* mRNA abundance (Fig. 4E), which was reflected in a reduced scaling of protein abundance with mRNA abundance across single cells (Fig 4F, G). To explore whether the suppressing effect of CSRP1 on *EEA1* translation acts through modulating the co-translational association of *EEA1* mRNA with early endosomes, we quantified the single-pixel correlations between *EEA1* smFISH signal and EEA1 antibody staining signal as well as the spatial patterns of *EEA1* transcripts. This showed a reduction in single-pixel intensity correlations, but no change in the distances of mRNA from the nuclear centroid and the spatial class assignment probabilities (Fig 4H, I). Taken together, these observations show that CSRP1 suppresses the translation of *EEA1* mRNA and may have an effect on *EEA1* mRNA localization to early endosomes. Whether these aspects are mechanistically coupled, and if this plays a role in linking *EEA1* translational efficiency to actin-mediated endocytosis and cellular state remains to be further explored.

In this study, we have reported the discovery of numerous mRNAs that associate with endosomes in a ribosome association-dependent manner, and we have explored some of the underlying mechanisms for the transcripts of *EEA1*, a well-known mediator of endosome tethering and homotypic fusion^20,21^. Amongst the mRNAs that associate to endosomes are multiple transcripts encoding for subunits of the WASH complex, a known regulator of the cortical actin network during epithelial morphogenesis as well as a direct regulator of endocytosis, endocytic recycling and retrograde transport, as well as multiple transcripts encoding proteins involved in growth factor and mTOR signaling. The WASH complex, together with the retromer complex, and actin also play a role in the organization of ER-endosome contact sites, and ER mediated endosomal fission^22,23,27^. Interestingly, we found that a subset of transcripts that encode for growth factor receptors, and which are translated on the ER, show a subcellular patterning that is similar to endosome-associated transcripts (see Fig. 1C). Possibly, this reflects a coordinated localization, and regulation of translation, of transcripts encoding for transmembrane proteins and cytoplasmic proteins with functions in the endo-lysosomal system at ER-endosome contact sites. This may also include transcripts encoding for cytoplasmic proteins involved in growth factor and mTOR signaling to couple this to cell proliferation and growth. Finally, we note that the localization of mRNAs on endosomes might play an important role in processes that are beyond the functioning of a single epithelial cell. Endosomes distribute asymmetrically in polarized cells, which plays important roles in cellular polarization (eg. brush border formation in enterocytes)^40^, in asymmetric cell division^41^, and in maintaining the stemness of pluripotent cells^42–44^. Combining multiplexed mRNA and protein readouts with RNA sequencing of organelle fractions and quantitative proteomics^17–19,24,45,46^ will be instrumental to study the role of subcellular transcript patterning in determining the phenotypic state of single cells and cell collectives, and their ability to display complex patterns of cellular decision making.

## Acknowledgements

We thank the members of Pelkmans lab for valuable discussions and René Holtackers for technical support. D.P. is supported by an EMBO (ALTF 1484-2015) and HFSP (LT-000531/2016) Long Term Fellowship. L.P. is supported by the Swiss National Science Foundation, the European Research Council (ERC Advanced Grant 885579), and the University of Zurich. W.N. and L.C.K. were supported by the Netherlands Organisation for Scientific Research (NWO; NWO-ALW-VENI 016.Veni.171.030 to W.N and NWO-ALW-VIDI 864.12.008 to L.C.K.) and the European Research Council (ERC Starting Grant 336291 to L.C.K. and ERC Consolidator Grant 819219 to L.C.K.).

## Author Contributions

L.P. and D.P. conceived the study; W.N. and L.K. invented the tools for chemical translocation of the organelles. W.N. generated the plasmids and cell lines for the chemical translocation of endosomes. D.P. performed all the experiments and analyzed the data. L.P. and D.P. wrote the manuscript.

**Figure S1.**
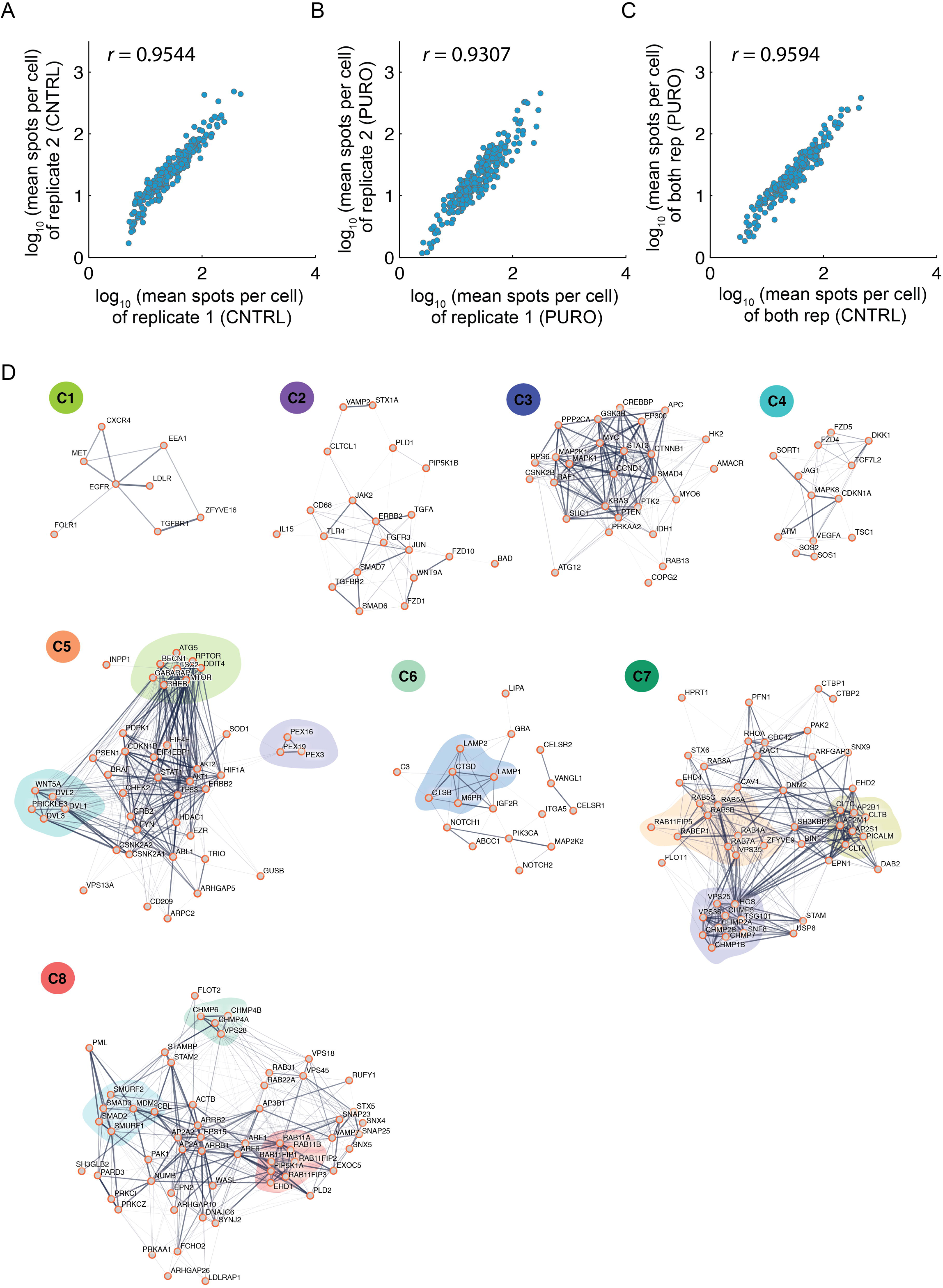
Image-based Screen for mRNAs that Localize on Endosomes. Related to Figure 1. (A) Pearson’s correlation of two replicates of measured mRNA count for each gene in Control condition. (B) Pearson’s correlation of two replicates of measured mRNA count for each gene in Puromycin treated condition. (C) Pearson’s correlation of mean spot count (for both Control replicates) in Control and mean spot count (for both Puromycin replicates) in Puromycin treated condition. (D) Networks of genes present within each cluster represented in Figure 1C. The networks were obtained from STRING, thickness of the line represents strength of their interaction (based on STRING).

**Figure S2.**
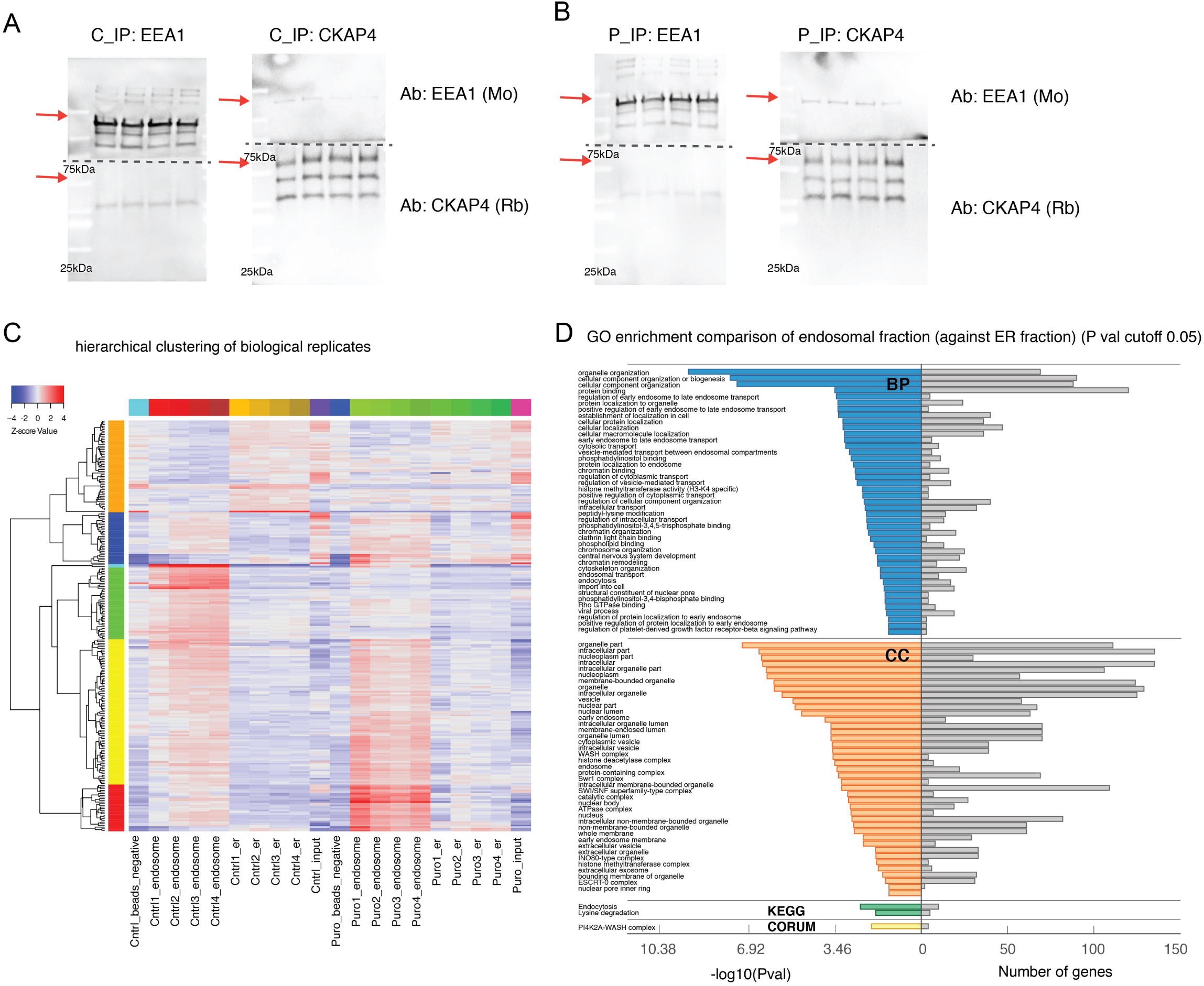
RNA sequencing of Purified Endosomes Reveals Associated Transcripts. Related to Figure 2. (A) Western blot representing 5% of total precipitated organelle fraction. Left side: precipitated endosomal control fraction, blotted against both markers: EEA1 (endosome); CKAP4 (ER). Right side: precipitated ER Control fraction, blotted against both markers: EEA1 (endosome); CKAP4 (ER). (B) Western blot representing 5% of total precipitated organelle fraction. Left side: precipitated endosomal puromycin fraction, blotted against both markers: EEA1 (endosome); CKAP4 (ER). Right side: precipitated ER puromycin fraction, blotted against both markers: EEA1 (endosome); CKAP4 (ER). (C) Hierarchical clustering of all genes in 4 biological replicates (Control and Puromycin), including empty beads controls and inputs. Expression values were Z-score normalized. (D) Gene Ontology enrichment analysis based on gProfiler, of all genes significantly present in endosomal Control fraction, compared to the ER Control fraction.

**Figure S3.**
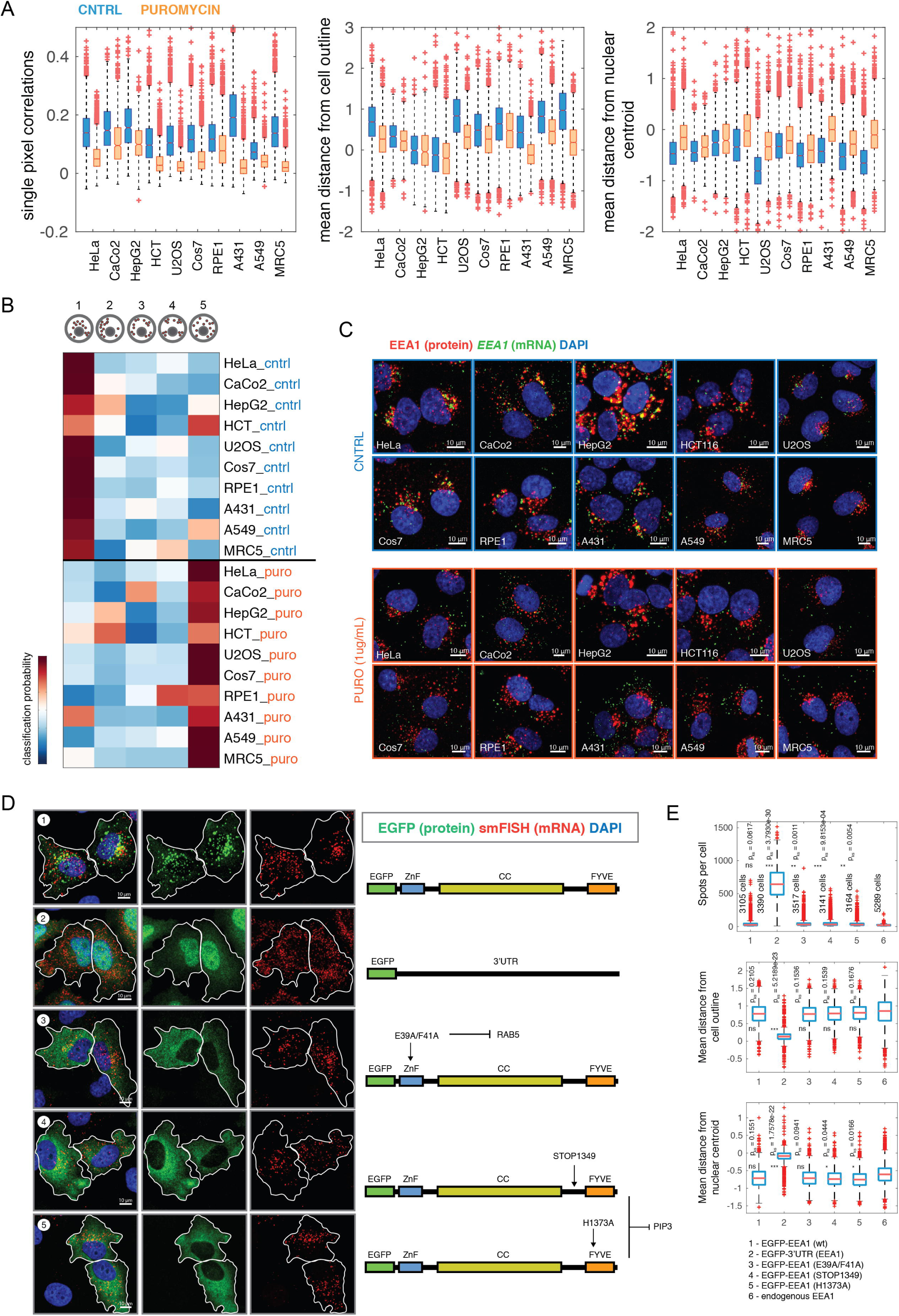
*EEA1* mRNA Localization is Conserved in Different Cell Lines and is Dependent on the cDNA Sequence. Related to Figure 3. (A) Left: boxplots representing single pixel correlations between sm-FISH signal and EEA1 protein stain in different cell lines, in control (blue) and puromycin (orange) treated condition. Middle: boxplots representing mean mRNA distance from the cell outline in different cell lines, in control (blue) and puromycin (orange) treated condition. Right: boxplots representing mean mRNA distance from the nuclear centroid in different cell lines, in control (blue) and puromycin (orange) treated condition. (B) Spatial classification probabilities of mRNA to adapt certain spatial pattern in different cell lines in control and puromycin treated condition. (C) Representative images of EEA1 mRNA and protein localization in different cells lines in control and puromycin treated condition. (D) HeLa cells expressing EGFP tagged variants of EEA1 cDNA and its mutated versions, or its 3’UTR region, 8 hours after induction of expression. sm-FISH was performed against EGFP sequence, and the green channel represents signal coming from EGFP (translated protein). (H) Boxplots of mRNA count of each EEA1 variant (as in (D)), its mean distance from the cell outline (middle plot) and nuclear centroid (lower plot). P values were calculated by performing KS test to compare the distributions of measured values across 100 bootstraps from the population of cells expressing different variants and those where endogenous EEA1 mRNA features were quantified.

**Figure S4.**
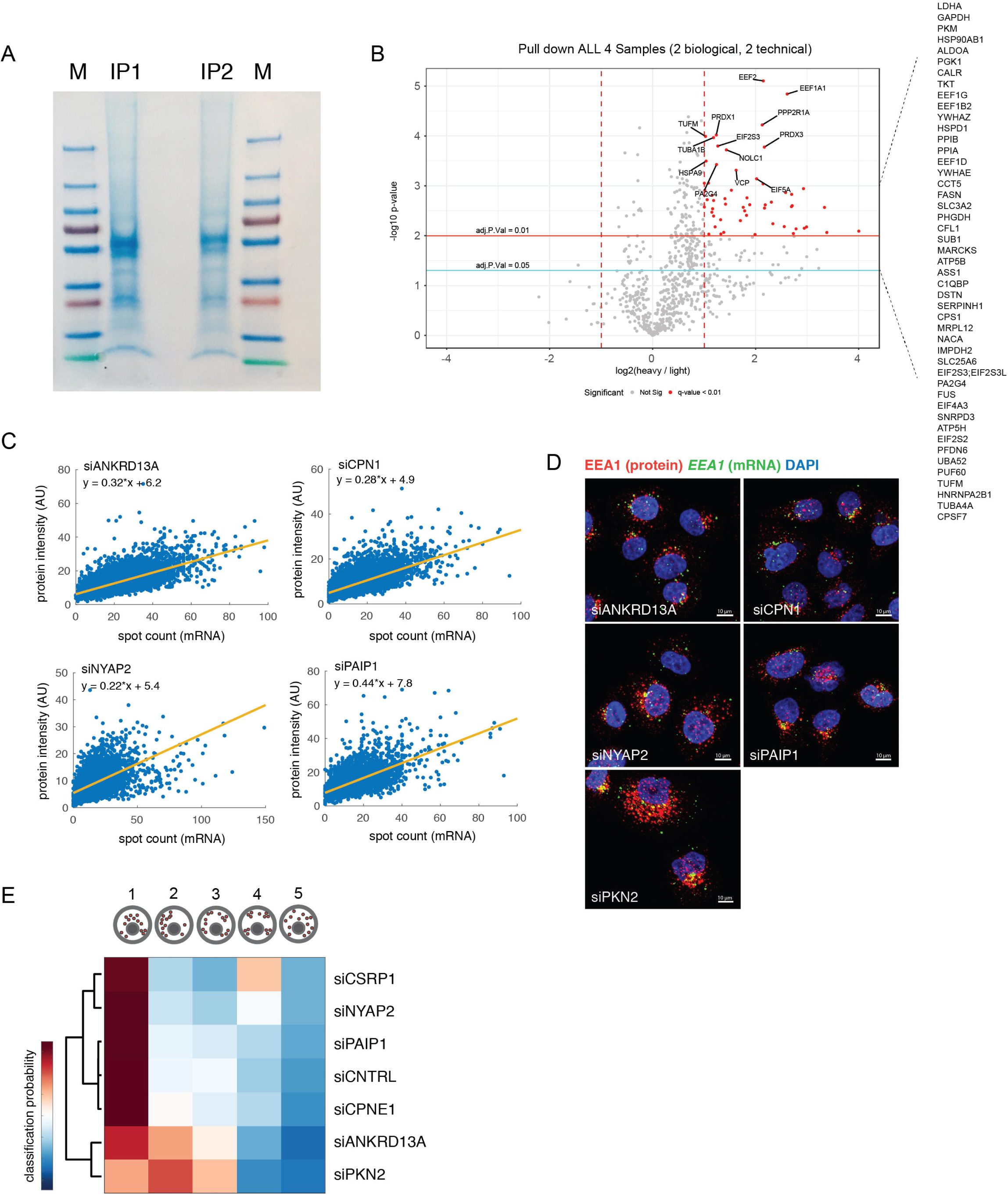
SILAC Based Proteomics Reveals Interactors of *EEA1* mRNA. Related to Figure 4. (A) Coomassie stained gel (10% of input) of precipitated lysates using magnetic oligo-coupled beads against *EEA1* mRNA (2 biological replicates). (B) Analysis of *ACTB* mRNA interactome (2 biological, 2 technical replicates). The most significant genes are listed on the right side of the volcano plot (pVal = 0.05). (C Linear regression plots using mRNA as a predictor for the protein quantities of EEA1 in cells treated with the siRNA against ANKRD13A, CPN1, NYAP2 and PAIP1. (D) Representative images of siRNA treated cells used for plotting in (C). Related to the Figure 4. (E) Spatial class probabilities of siCNTRL and siRNA treated cells (against *EEA1* mRNA interactors).

## Contact for Reagent and Resource Sharing

Further information and requests for resources and reagents should be directed to and will be fulfilled by the Lead Contact

## Experimental Model and Subject Details

### Cell Lines Source and Cultivation

HeLa cell lines (Kyoto) were obtained from Jan Ellenberg (EMBL, Heidelberg, Germany). All other cell lines were previously obtained from ATCC and cultured within the lab, including HEK 293T cell lines used for the production of Lentivirus.

## Method Details

### Experimental Design Details

Datasets represented in the figures for all cell measurements contain two biological replicates, unless stated otherwise.

### Image-based Transcriptomics and Antibody Labelling

Image-based transcriptomics and image processing was performed using open source software CellProfiler and our custom made Matlab modules available at, and https://github.com/pelkmanslab/ImageBasedTranscriptomics/tree/master/CellProfiler/Modules. Briefly, the modules have been at first tested locally on several images and obtained segmentation images manually inspected. Upon evaluating the good segmentation, pipeline was run on the whole image dataset, using ScienceCloud computational infrastructure of University of Zurich (UZH), or computational cluster Brutus (ETH).

For all cell lines final dilution of the protease was 1:16000. Protease was first diluted 1:8000 in 1xPBS and dispensed in volume of 30uL, prior to which the volume within the well was aspirated to 30uL, such as is recommended in the protocol for bDNA smFISH (ThermoScientific, previously Affymetrix). Cells were seeded in 96-well plates. Image-based transcriptomics screen was performed in 384-well plate format. Branched DNA single-molecule fluorescence *in-situ* hybridization was performed using ViewRNA reagents (Affymetrix, Thermo Fisher Scientific). All wash steps were performed on the Biomek robotic liquid handler. Imaging was done using automated confocal microscope, CellVoyager 7000 (Yokogawa) with the enhanced CSU-X1 spinning disc (Microlens enhanced dual Nipkow disc confocal scanner, wide view type) and a 40X Olympus objective of 0.95 NA and Neo sCMOS cameras (Andor, 2.560 × 2.560 pixels). 10 Z slices with the distance of 1uM were imaged and maximum projection images (MIP) used for the further image analysis.

### Plasmids and cloning

CSRP1-GFP-Spark was purchased from Sino Biological (cat nr. HG11201-AG). EEA1 plasmids were generated using Gateway cloning system (Thermo Fisher Scientific), and EGFP-EEA1 plasmid as a template for cloning (Addgene: #42307). Full length EEA1 was introduced in pEGFP-Blast destination vector (Wade Harper lab), after being subcloned first to pENTR223. All point mutants of EEA1 were generated by site directed mutagenesis on pENTR223 vector, and subsequently introduced to the destination vector.

pMD2.G (Addgene#12259) and psPAX2 (Addgene #12260) were gifts from Didier Trono. pSIN-TRE-rtTA-IRES self-inactivating lentiviral vectors were constructed from pSIN-TRE-rtTA-IRES-Puro (a gift from Benjamin Bouchet, Utrecht University, The Netherlands), by replacing the puromycin-resistance cassette with either the hygromycin-resistance cassette from pCDNA5-FRT-TO (Invitrogen) to generate pSIN-TRE-rtTA-IRES-Hygro, or with the blasticidin-resistance cassette from pBabe-blast (a gift from Geert Kops, Hubrecht Institute-KNAW, The Netherlands) to generate pSIN-

TRE-rtTA-IRES-Blast. pSIN-TRE-rtTA-IRES-Blast-KIF1A(1-365)-FLAG-FRB encodes amino acids 1-365 of murine KifA (derived from pB80-KIF1A(1-365)-GFP-SSPB(micro)^47^, C-terminally fused to a 2xFLAG tag (DYKDHDGDYKDHD) and the FKBP-rapalog-binding domain FRB (derived from FRB-TagBFP-linker-LOVpep^30^. pSIN-TRE-rtTA-IRES-Hygro-FKBP-mCherry-RAB5, -RAB7 and -RAB11 encode the full length wildtype version of either human RAB5, decorating early endosomes (derived from pB80-iLID-mCherry-RAB5^47^, human RAB7, decorating late endosomes (derived from GW1-GFP-Rab7, a gift from Casper Hoogenraad, Utrecht University, The Netherlands) or human RAB11 (derived from pB80-iLID-mCherry-RAB11^47^, fused N-terminally to the rapalog acceptor FKBP and mCherry (derived from pB80-FKBP-mCherry-RAB11^47^. All constructs were validated by sequencing of the full ORF.

### Transient Transfection of cDNA

HeLa cells were seeded in 96-well plate and cultivated for 36h prior to the transfection. cDNA coding for CSRP1-GFP-Spark was transfected using GeneJuice reagent (Novagen) according to the manufacturer protocol and grown for another 24 hours. Cells were subsequently fixed and stained.

### Lentiviral transduction

Lentivirus packaging was performed by using MaxPEI-based co-transfection of HEK293T cells with psPAX2, pMD2.G and the pSIN-TRE-rtTA-IRES lentiviral vectors. Supernatant of packaging cells was harvested up to 72 h after transfection, filtered through a 0.45-µm filter and incubated with a polyethylene glycol (PEG)-6000-based precipitation solution overnight at 4°C. After precipitation, virus was concentrated up to 100× by centrifugation and dissolution in 1× phosphate buffered saline (PBS).

HeLa Kyoto cells were incubated for 1 h in full growth medium supplemented with 8 µg/ml polybrene before infection with lentivirus encoding doxycycline-inducible KIF1A(1-365)-FLAG-FRB and FBKP-mCherry-RAB5, 7 or 11. To establish clonal stable lines, medium was supplemented with μg/ml hygromycin and 10 μg/ml blasticidin (both from Invivogen) at 48 hrs after infection. Subsequently, cells were grown to confluency and seeded as single cells. Single clones were selected to have correct labeling of FKBP-mCherry-RAB protein and to display rapalog-induced organelle repositioning by live cell microscopy. Doxycycline-inducible expression of KIF1A(1-365)-FLAG-FRB and FKBP-mCherry-RAB protein was validated by immunoblotting.

### Induced endosome repositioning

To couple FKBP-mCherry-RAB to KIF1A(1-365)-FLAG-FRB, rapalog (AP21967, ARIAD) dissolved in ethanol was added to the cells to establish a final rapalog concentration of 100nM. Cells were incubated together with the rapalog containing medium for 45 minutes, subsequently fixed with 4% PFA and subjected to the smFISH protocol.

## Quantification and Statistical Analysis

### Feature Extraction

Area, shape, intensities and texture (at a scale of 5 pixels) of cells and nuclei were extracted using open source software CellProfiler. Correction of uneven illumination and subtraction of camera dependent invariant background was performed using custom CellProfiler modules as previously described. Briefly, large number of acquired images per channel can be used to learn pixel-wise illumination and signal gain biases. For each pixel, standard deviation and mean intensity value is calculated for a given channel. Illumination bias is than corrected by performing Z-scoring per pixel and reverting the values to the intensity values. Population context features were measured using previously published module implemented within the iBRAIN pipeline that calculates Local Cell Density and Distance to Edge for each single cell, after completion of the CellProfiler pipeline. Code for generation of population context features can be found on our Github repository https://github.com/pelkmanslab/iBRAINShared/tree/master/iBRAIN/CreatePopulationContext. Number of directly adjacent cells and size of extracellular space with overlap to other cells was extracted using custom CellProfiler module, extending the cell outline by 10 pixels.

Cells that had multiple nuclei, border cells and missegmented cells were discarded using supervised machine learning (SVM) tool CellClassifier, available at https://www.pelkmanslab.org/?page_id=63. Briefly, images with overlayed segmentations of cells and their nuclei have been loaded in Matlab GUI of CellClassifier and classifier was manually trained by selecting cells with wrongly segmented nucleus as a class 1, and correctly segmented nucleus as class 2. Subsequently classifier was tested on a randomly selected images and applied to the whole dataset. Features of nuclear shape and DAPI intensity were used for the classification. Similarly, features of cellular shape were used to train classifier and recognize correctly segmented cells.

### Selection of Genes for the Image-Based Transcriptomic Screen

Image-based screen was performed on targeted subset of genes for which bDNA sm-FISH probes were available in our previously published library. The gene selection was made based on the functional classification of genes, so that primarily those involved either in endocytosis and endocytic signaling and endomembrane trafficking and regulation are inspected. This resulted in the subset of 328 probe sets.

### Isolation of Endosomal and ER Fractions Using Magnetic Beads

To isolate and purify the fractions of intracellular organelles, we have applied homogenization protocol, usually performed prior to the standard organelle fractionation using differential centrifugation on sucrose gradient. Specifically, HeLa cells were first grown in 15cm dishes, to the confluency of 90%. Cells were first washed using 1xPBS while still being attached to the bottom of the dish. Subsequently, cells were scrapped of the dish using cell scrapper, and 1xPBS, and span down on 1000G for 5 min. Cellular pellet was than washed gently by up and down pipetting in 2mL of sucrose-based homogenization buffer (4% sucrose, 3mM imidazole, pH 7.4, + Roche tablet EDTA free inhibitors, +Roche Phosphostop tablet), and span down on 3000G for 5 min. Buffer was aspirated, and new homogenization buffer added on cells (each pellet dissolved in 1mL of buffer), and cells were than homogenized by passing the suspension 12 times through the Isobiotec Cell homogenizer (18 micron). Next, 5 microliters of cell homogenate were at first carefully inspected on the microscope, to reassure efficient release of intracellular organelles, and upon confirmation of efficient lysis, lysate was span down on 3000G for 10 minutes. Supernatant was then used for the purification of organelles, such that it was transferred to the prewashed, antibody coupled magnetic beads (Dynabeads), gently mixed, and incubated at 4°C overnight. Next morning, beads were washed first three times using sucrose-based homogenization buffer, followed by one wash in 1xPBS. Subsequently, such washed beads were incubated in buffers from Qiagen RNA isolation Kit, and processed similarly as cellular lysates, so that the all RNA bound to the beads could be purified using Quiagen RNA affinity column. Total RNA was eluted in 30 µL of the ddH_2_O, and subsequently used for the quality control check, generation of library and NGS sequencing.

### Illumina RNA Sequencing Experiment Description

#### Library Preparation

The quality of the isolated RNA was determined with a Qubit® (1.0) Fluorometer (Life Technologies, California, USA) and a Bioanalyzer 2100 (Agilent, Waldbronn, Germany). Only those samples with a 260 nm/280 nm ratio between 1.8–2.1 and a 28S/18S ratio within 1.5–2 were further processed. The TruSeq RNA Sample Prep Kit v2 (Illumina, Inc, California, USA) was used in the succeeding steps. Briefly, total RNA samples (100-1000 ng) were ribo-depleted using Ribo Zero Gold (Epicentre®, USA) and then fragmented. The fragmented samples were reversed transcribed to cDNA, end-repaired and polyadenylated before ligation of TruSeq adapters containing the index for multiplexing Fragments containing TruSeq adapters on both ends were selectively enriched with PCR. The quality and quantity of the enriched libraries were validated using Qubit® (1.0) Fluorometer and the Caliper GX LabChip® GX (Caliper Life Sciences, Inc., USA). The product is a smear with an average fragment size of approximately 260 bp. The libraries were normalized to 10nM in Tris-Cl 10 mM, pH8.5 with 0.1% Tween 20.

#### Cluster Generation and Sequencing

The TruSeq PE Cluster Kit v4-cBot-HS or TruSeq SR Cluster Kit v4-cBot-HS (Illumina, Inc, California, USA) was used for cluster generation using 10 pM of pooled normalized libraries on the cBOT. Sequencing were performed on the Illumina HiSeq 2500 paired end at 2 X101 bp or single end 100 bp using the TruSeq SBS Kit v4-HS (Illumina, Inc, California, USA).

### Template RNA-Seq Data Analysis

The RNA-seq data analysis consisted of the following steps:

1. The raw reads were first cleaned by removing adapter sequences, trimming low quality ends, and filtering reads with low quality (phred quality <20) using Trimmomatic (Version 0.36) [Trimmomatic].
2. The read alignment was done with STAR (v2.5.3a) [STAR]. As reference we used the Ensembl genome build GRCh38.p10 with the gene annotations downloaded on 2018-02-26 from Ensembl (release 91). The STAR alignment options were “-- outFilterType BySJout --outFilterMatchNmin 30 --outFilterMismatchNmax 10 -- outFilterMismatchNoverLmax 0.05 --alignSJDBoverhangMin 1 –alignSJoverhangMin 8 --alignIntronMax 1000000 --alignMatesGapMax 1000000 –outFilterMultimapNmax 50”.
3. Gene expression values were computed with the function featureCounts from the R package Rsubread (v1.26.0) [Rsubread]. The options for featureCounts were: - min mapping quality 10 - min feature overlap 10bp - count multi-mapping reads - count only primary alignments - count reads also if they overlap multiple genes.
4. To detect differentially expressed genes we applied a count based negative binomial model implemented in the software package EdgeR (R version: 3.5.0, EdgeR version: 3.22.1) [EdgeR]. The differential expression was assessed using an exact test adapted for over-dispersed data. Genes showing altered expression with adjusted (Benjamini and Hochberg method) p-value < 0.05 were considered differentially expressed.

### SILAC Based Proteomics of *ACTB* and *EEA1* mRNA

Stable Isotope Labeling by Amino acids in Culture was performed on HeLa cells using following reagents: Lysine-stock solutions: for K0: L-Lysine (L8662, Sigma), for K8: L-Lysine, 2HCl U-13C U15N (CNLM-291, Cambridge Isotope Laboratories). Lysine was dissolved in 1xPBS to a final concentration of 146mg/ml, filtered through 0.22 μm filter, and stored at 4°C and in the dark (aluminum foil wrapped). Arginine-stock solutions: for R0: L-Arginine (A8094, Sigma), for R10: L-Arginine, HCl U-13C6 U-15N4 (CNLM-539, Cambridge Isotope Laboratories). Arginine was dissolved in 1xPBS to a final concentration of 84mg/ml and filtered through 0.22 μm filter and stored at 4°C and in the dark (aluminum foil wrapped). SILAC media was prepared using SILAC DMEM (Life Technologies, #A14431-01), add 5 ml 100x L-Glutamine, 5 ml 100x Pen/Strep and 5ml 100mM Sodium Pyruvate (Gibco, #11360-039), 250 μl Arginine-Stock (84 mg/ml) and 250 μl Lysine-Stock (84 mg/ml) and filtered through 0.22 μm 500 ml filter system. On each 500 ml of media 50 ml of dialyzed FBS (GIBCO, #26400-044) was added. Each precipitation was performed on lysate originating from 3 15cm dishes of HeLa cells, that were washed in 1xPBS and subsequently lysed in lysis buffer (50mM Tris/HCl, 150mM NaCl, 1mM MgCl2, pH 7.4) supplemented with the Roche tablet of complete inhibitors without EDTA and Phosphostop tablet (Roche). Beads used for the precipitation of the mRNA were obtain from Affymetrix (Thermo Fisher Scientific), custom made, coupled to the oligos against sequence of human *ACTB, EEA1* or *EGFP* respectively. Lysates were incubated with the beads over night at 4°C, and beads were washed three times with the lysis buffer, followed by one wash in 1xPBS and last wash in ddH_2_O, leaving 20 μl of total volume, from which 2 μl were taken and ran on the gel prior to the processing for further MS/MS analysis.

### Sample preparation

Samples were prepared using a commercial iST Kit (PreOmics) with an updated version of the protocol. Briefly, the washed beads of each sample were re-suspended in 50 µl ‘Lyse’ buffer and incubated for 10 min at 95°C. 100 µg of each sample was transferred to the cartridge and digested by adding 50 µl of the ‘Digest’ solution. After 60 min of incubation at 37°C the digestion was stopped with 100 µl of ‘Stop’ solution. The solutions in the cartridge were removed by centrifugation at 3800xg, while the peptides were retained by the iST-filter. Finally, the peptides were washed, eluted, dried and re-solubilized in 15 µl ‘LC-Load’ solvent for MS-Analysis.

### Mass spectrometry

Dissolved samples were injected by an Easy-nLC 1000 system (Thermo Scientific) and separated on an EasySpray-column (75 µm x 500 mm) packed with C18 material (PepMap, C18, 100 Å, 2 µm 50°C, Thermo Scientific). The column was equilibrated with 100% solvent A (0.1% formic acid (FA) in water). Peptides were eluted using the following gradient of solvent B (0.1% FA in ACN): 5% B for 2min; 5-25% B in, 90 min; 25-35% B in 10 min; 35-99% B in 5 min at a flow rate of 0.3 µl/min. High accuracy mass spectra were acquired with an Orbitrap Fusion (Thermo Scientific) that was operated in data depended acquisition mode. All precursor signals were recorded in the Orbitrap using quadrupole transmission in the mass range of 400-1500 m/z. Spectra were recorded with a resolution of 120 000 at 200 m/z, a target value of 5E5 and the maximum cycle time was set to 3 seconds. Data dependent MS/MS were recorded in the linear ion trap using quadrupole isolation with a window of 1.6 Da and HCD fragmentation with 32% fragmentation energy. The ion trap was operated in rapid scan mode with a target value of 1E4 and a maximum injection time of 50 ms. Precursor signals were selected for fragmentation with a charge state from +2 to +7 and a signal intensity of at least 5E3. A dynamic exclusion list was used for 30 seconds and maximum parallelizing ion injections was activated. The mass spectrometry proteomics data were handled using the local laboratory information management system (LIMS)^48^ and all relevant data have been deposited to the ProteomeXchange Consortium via the PRIDE (http://www.ebi.ac.uk/pride) partner repository with the data set identifier PXDXXXX.

### Protein identification and label free protein quantification

The acquired raw MS data were processed by MaxQuant (version 1.6.2.10), followed by protein identification using the integrated Andromeda search engine^49^. Spectra were searched against the Human database UniProtKB (release 20180405, 20336 entries) concatenated to common protein contaminants. Carbamidomethylation of cysteine was set as fixed modification, while methionine oxidation and N-terminal protein acetylation were set as variable. Enzyme specificity was set to trypsin/P allowing a minimal peptide length of 6 amino acids and a maximum of one missed cleavage. The maximum false discovery rate (FDR) was set to 0.01 for peptides and 0.01 for proteins. Lys8 and Arg10 were used as heavy labels and a 0.7 min window for match between runs was applied. Heavy to light ratios of the protein group result were analyzed further with Perseus (version 1.6.2.2)^50^.

### Comparison of Distributions using Kolmogorov-Smirnov Statistics

The distributions of the replicates or control and perturbed datasets were compared using Matlab function “kstest2”. We performed 100 bootstraps over all cells, calculated p values across all subsampling and represented mean p value across all bootstraps as a final (within the Figures).

### Classification of Cells Based on the Transcript Spatial Pattern

Probabilities to belong to a specific spatial class was calculated based on the set of spatial features extracted using our custom CellProfiler module. Briefly, we first calculated the primary set of features using module MeasureLocalizationOfSpots.m that is available on our Github. Subsequently, for every cell and set of primary spatial features, cellular features were obtained by the custom MeasureChildren.m CellProfiler module, available at https://github.com/pelkmanslab/ImageBasedTranscriptomics/tree/master/CellProfiler/Modules. Further, only cellular features describing mean and sd of spatial features for each single cell were used to perform hierarchical clustering of cells using Euclidean distance space and Ward’s linkage method. Cells were grouped in 50 bins, and 1000 samplings of 100 cells was performed using script available at https://github.com/pelkmanslab/locpatterns. To classify clusters from those different samplings into spatial pattern types the centroid for each cluster was computed (using 7 centroids), and distance to a number of randomly sampled cluster centroids measured. The localization type of the cell was than defined as that of its closest centroid, and probability as a fraction of times a cell was defined to belong to a particular localization type.

### siRNA Knock-down Experiments

siRNA was performed by transfection of two different oligos against gene of interest synthesized by Microsynth and transfected in final concentration 10 nM each (total 20nm). Cells were reverse transfected, by seeding in 96 well plate (Greiner) that already contained mix of RNAi-Max reagent and siRNA, in OptiMEM. 1000 cells per well was seeded at the time of transfection. Cells were incubated together with the siRNA mix overnight, and washed next morning using Biomek wash dispenser with full serum containing media (DMEM, 10% FBS) and left for next 72 hours to grow in the incubator.

siRNA Oligo sequences (as designed based on Dharmacon):

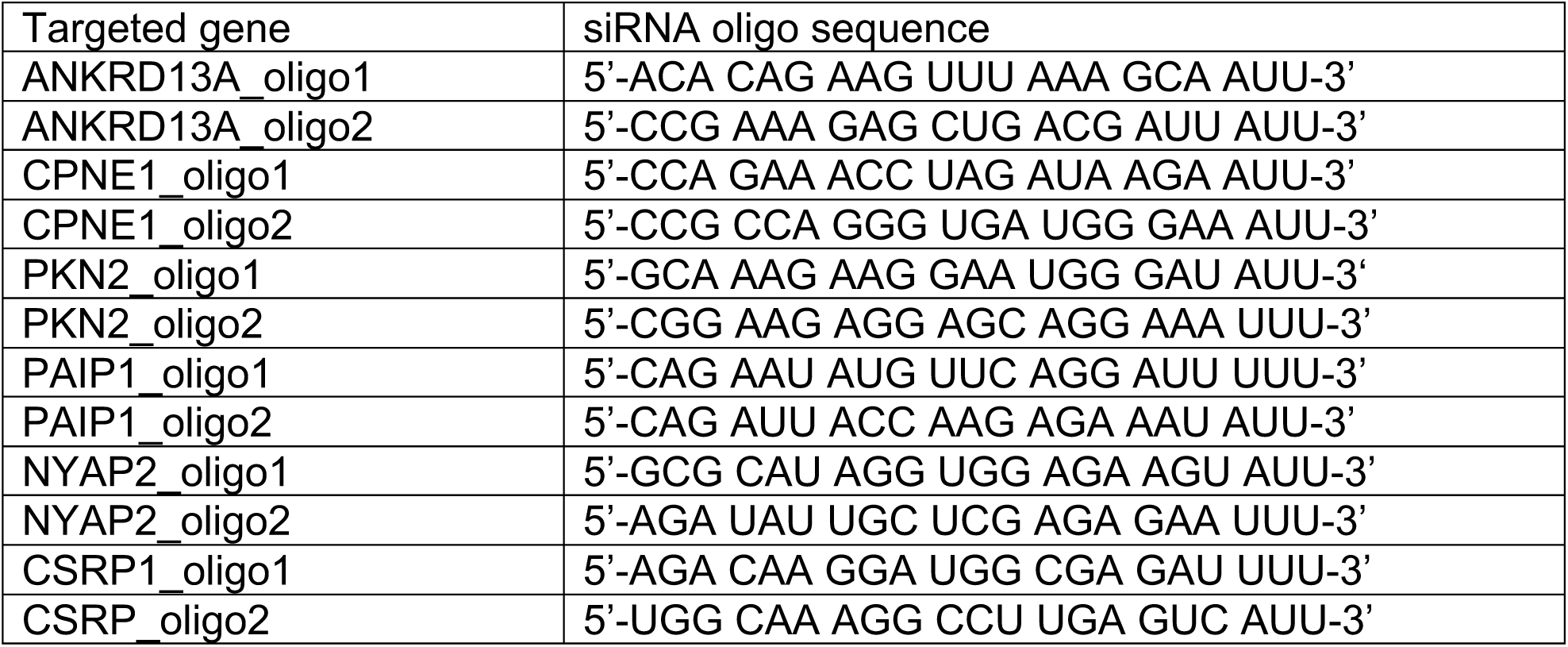

### CRISPR/Cas9 Knock-out Experiments and Rescue

Knock-out of EEA1 was achieved by transient transfection of plasmid containing sequence for CRISPR/Cas9 mediated KO of EEA1 (gRNA sequence: TTACCTTAAGTGAAGCCTGT) ^50^. Subsequently, cells were seeded in dilution of less than 1 cell per well, in plastic 96 well plate, and left to grow for 4-5 weeks. Upon expansion of single clones, cell colonies were inspected for expression of EEA1 by sm-FISH and immunolabeling and single clones with preserved cellular morphology and cell cycle length were used for reconstitution experiments, using Gateway destination plasmids encoding either full length or mutant version of EGFP tagged EEA1.

